# CircTulp4 functions in Alzheimer’s disease pathogenesis by regulating its parental gene, Tulp4

**DOI:** 10.1101/2020.07.08.192716

**Authors:** Nana Ma, Jie Pan, Qi Wu, Bo Yu, Jun Wan, Wei Zhang

## Abstract

Alzheimer’s disease (AD)—one of the most common neurodegenerative diseases worldwide—impairs cognition, memory, and language ability and causes dementia. However, AD pathogenesis remains poorly elucidated. Recently, a potential link between AD and circular RNAs (circRNAs) has been uncovered, but only a few circRNAs that might be involved in AD have been identified. Here, we systematically investigated circRNAs in the APP/PS1 model mouse brain through deep RNA-sequencing. We report that circRNAs are markedly enriched in the brain and that several circRNAs exhibit differential expression between wild-type and APP/PS1 mice. We characterized one abundant circRNA, circTulp4, derived from Intron1 of the gene *Tulp4*. Our results indicate that circTulp4 predominantly localizes in the nucleus and interacts with U1 snRNP and RNA polymerase II to modulate the transcription of its parental gene, *Tulp4*, and thereby regulate the function of the nervous system and might participate in the development of AD.

## Background

Only ~2% of the genes in the human genome encode proteins [1]. Thus, non-protein-coding RNAs such as circular RNAs (circRNAs) and microRNAs (miRNAs/miRs) are now under considerable research scrutiny and generally accepted to influence myriad physiological and pathological processes, including the initiation and progression of neurological diseases such as Alzheimer’s disease (AD) [2–6], which is associated with progressive loss of memory, cognition, and behavioral capabilities [7,8]. No effective AD treatments are available, and the genetic basis of AD and the epigenetic control of AD pathogenesis remain largely unelucidated.

CircRNAs are unique among cellular RNAs in that pre-mRNA back-splicing results in covalent linking of the 5′- and 3′-ends [9], which is likely responsible for circRNAs being more stable than linear coding/noncoding transcripts [10]. Given this high stability (lifetimes of hours to days or longer), circRNAs could perform functions distinct from those of linear RNAs and also potentially serve as more reliable biomarkers of health and disease than other RNAs. Intriguingly, in the brain, circRNAs are enriched in neurons and in synaptosomes [11] and their levels are regulated during aging in the fly and the mouse; thus, these circRNAs might play roles in brain aging and aging-associated neurodegenerative diseases [12,13]. For example, CDR1as (CiRS-7: circRNA sponge bound to miR-7) and miR-7 are associated with nervous system development and disease, and CDR1as dysfunction can cause miR-7 upregulation and lead to downregulation of AD-related targets, including ubiquitin-protein ligase A [14–16]. Furthermore, an *SRY*-derived circRNA can act as a natural miRNA sponge to inhibit miR-138 activity [17], which can affect learning and memory by regulating acyl protein thioesterase 1 [18,19]. However, circRNAs and their activities in AD have not been comprehensively analyzed.

To identify AD-associated circRNAs, we used deep RNA-sequencing (RNA-seq) for the first comparative profiling of circRNA expression in the brain of APP/PS1 mice (which express APP_695swe_ and carry PS1-dE9 mutations) and wild-type (WT) mice. We used 6- and 9-month-old mice because β-amyloid appears at 6 months and extracellular β-amyloid deposits in the cortex are apparent by 9 months, and synaptic transmission and long-term potentiation are impaired when the mice are 9–12 months old [20]. Our circRNA-profiling data could serve as a useful resource for future development of AD therapeutic targets or novel diagnostics. Moreover, our findings identify circTulp4 as a potential AD biomarker: circTulp4 is enriched in the brain and downregulated in APP/PS1 mice, and circTulp4 mainly localizes in the nucleus, interacts with U1 snRNP and RNA polymerase II to promote the transcription of its parental gene, *Tulp4*, and by affecting *Tulp4* expression, influences neuronal growth and differentiation.

## Results

### Brain circRNA profiles differ in WT and APP/PS1 mice

We performed RNA-seq analyses on rRNA-/linear-RNA-depleted total RNA isolated from the brain of 6-/9-month-old WT and APP/PS1 mice; from the final clean reads, we identified 15,713 circRNAs (See Supplementary Data). The circRNA candidates were annotated using RefSeq database [22]. Most circRNAs were <5000 nt long (median length, 1000 nt; Fig. 1A), and the majority aligned with protein-coding exons and fewer with introns and intergenic regions (Fig. 1B). We normalized the junction reads (supporting the identification of RNAs as circRNAs) by read length and number of mapped reads (spliced reads per billion mapped reads, denoted as TPM), which permitted quantitative comparisons between junction reads from different RNA-seq data (Fig. 1C). Numerous circRNAs exhibited differential expression across groups, which we quantified using Wilcoxon rank-sum test. In 6-month-old mice, 343 circRNAs were differentially expressed between the two groups: 192 and 151 transcripts were significantly (p<0.05) upregulated and downregulated, respectively, in APP/PS1 mice relative to WT (Fig. 1D); at 9 months, 243 circRNAs were significantly dysregulated: 141 and 102 transcripts were upregulated and downregulated in APP/PS1 mice (Fig. 1E). To generate these volcano maps, cluster analysis was performed on the differentially expressed circRNAs. The brain circRNA profiles of APP/PS1 and WT mice differed (Fig. 1F), and the parental genes of the upregulated/downregulated circRNAs mapped to distinct chromosomes (Fig. 1G).

**Figure 1.**
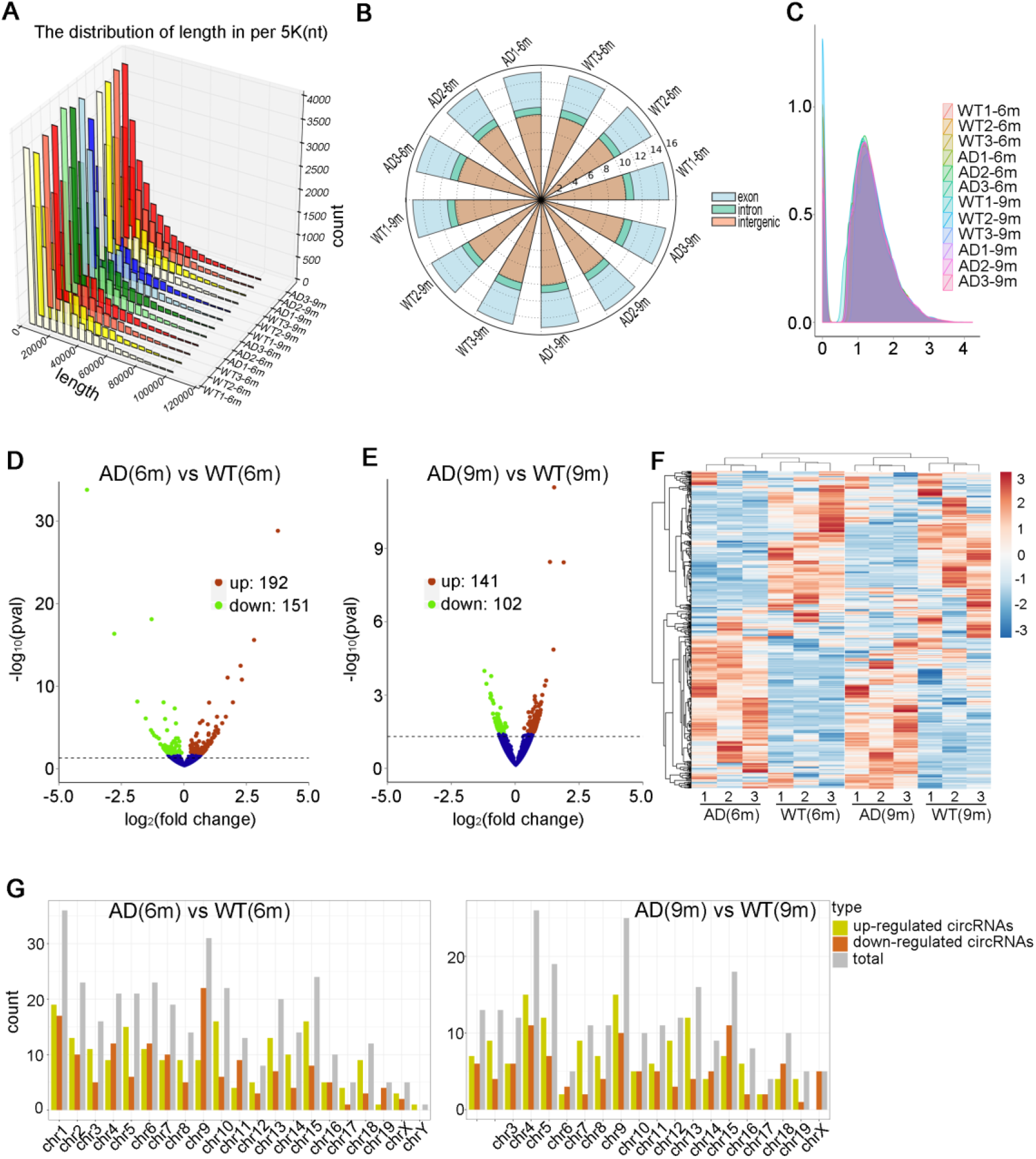
CircRNA profiling in the brain of WT and APP/PS1 mice. **a** CircRNA length distribution: x-axis: lengths of circRNAs detected in this study; y-axis: abundance of circRNAs classified by lengths; z-axis: sample information. **b** CircRNAs derived from distinct genomic regions; circRNA numbers are listed as log2N, where N is the number of exonic, intronic, or intergenic reads. For example, from the origin, the number of exonic reads is ~2^15^. **c** Density plot of circRNA abundance in WT and APP/PS1 mice. **d, e** Volcano plots showing circRNAs that exhibited downregulation (green points), upregulation (red points), or no statistically significant difference (blue points) in WT and APP/PS1 mice; x-axis: log2 ratio of circRNA expression levels between WT and APP/PS1 mice; y-axis: false-discovery rate (-log10 transformed) of circRNAs. **f** Heatmap of all circRNAs differentially expressed between WT and APP/PS1 mouse brains. **g** Numbers of differentially expressed circRNAs in different chromosomes; yellow, red, and gray bars: upregulated, downregulated, and total circRNAs, respectively.

We next verified that back-splicing events indicated the genuine circular form of the RNA transcripts and further confirmed the differential expression identified using RNA-seq data. For this real-time qPCR analysis, we designed divergent primers against 26 circRNAs (Supplementary Fig. 1); each primer pair amplified a single product of the expected size. All transcripts were detected in the brain of 2–9-month-old WT and APP/PS1 mice and exhibited significant differential expression. Our qPCR results were highly consistent with the RNA-seq data.

### CircTulp4 is a circRNA that mainly localizes in the nucleus and regulates its parental gene

Our RNA-seq screening identified an array of brain circRNAs showing altered expression in APP/PS1 mice relative to WT. We now focus on our investigation on one specific circRNA—circTulp4—as a potential AD biomarker. CircTulp4, derived from *Tulp4* Intron1 (Fig. 2A), was reported to be markedly enriched in the mammalian brain and to feature a sequence well conserved in humans and mice [11]. Our high-throughput sequencing data revealed high circTulp4 levels in the brain, and our qPCR results confirmed significant differential expression of circTulp4 between APP/PS1 and WT mice at both 9 and 12 months (Fig. 5A).

**Figure 2.**
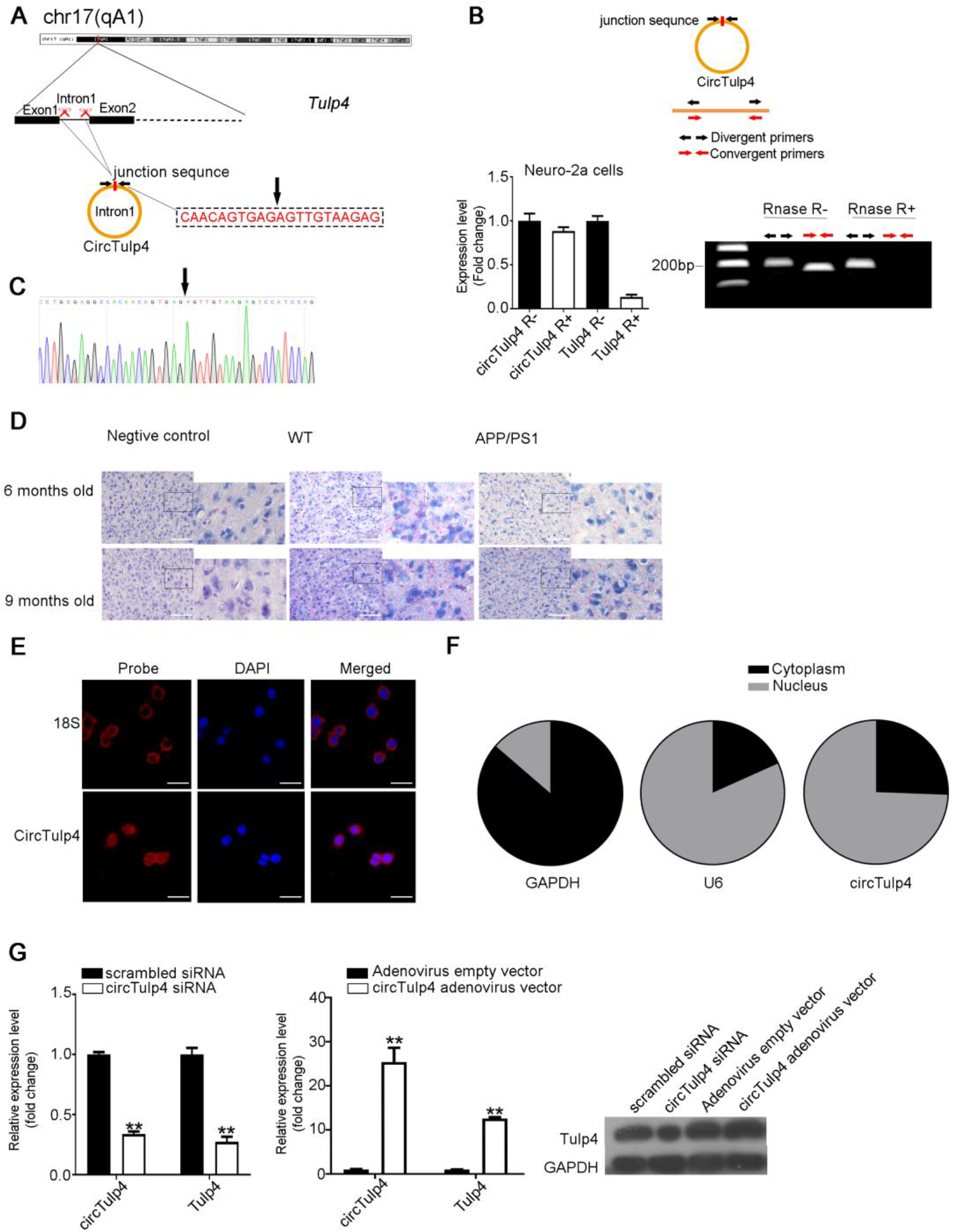
CircTulp4 is a circRNA that mainly localizes in the nucleus and regulates its parental gene, *Tulp4*. **a** Schematic depicting circTulp4 formation through circularization of Intron1 in *Tulp4*. **b** Upper panel: Convergent (red) and divergent (black) primers designed to amplify linear and back-spliced products. Lower panel: qRT-PCR analysis of abundance of circTulp4 and *Tulp4* mRNA in Neuro-2a cells treated with RNase R; circTulp4 and *Tulp4* mRNA amounts were normalized to the value measured in mock treatment. **c** Sanger sequencing after PCR performed using the indicated divergent flanking primers, confirming “head-to-tail” splicing of circTulp4 in Neuro-2a cells. **d** CircTulp4 expression detected in fresh-frozen brain tissue samples from 6- and 9-month-old WT and APP/PS1 mice, using BaseScope® 2.5 HD visible light staining kit; circTulp4 appears as red dots. Cells were counterstained with hematoxylin. Single-pair probes for circTulp4 targeting a unique junction sequence are specific and sensitive. Original magnification 40×; scale bar : 110 μm. **e** RNA fluorescence in situ hybridization (FISH) for circTulp4; 18S is a cytoplasmic control. Nuclei were stained with 4,6-diamidino-2-phenylindole (DAPI); circTulp4 and 18S probes were labeled with Alexa Fluor 555. Original magnification 40×; scale bar: 40 μm. **f** qRT-PCR data indicating circTulp4 abundance in cytoplasm and nucleus of primary hippocampal neurons, determined by using nuclear mass-separation assay; GAPDH and U6 are shown for comparison. **g** Tulp4 mRNA and protein levels were decreased after siRNA-mediated knockdown of circTulp4 in primary hippocampal neurons; the specific siRNA targeted circTulp4 at junction sequences. Results for control mismatched-sequence siRNA are shown in black. Parental-gene mRNA and protein levels were increased in primary hippocampal neurons after overexpression of circTulp4 by using a circTulp4 adenovirus vector; adenovirus empty-vector results are shown in black. Results shown are means ± SD (*p<0.05, **p<0.01).

To verify that *Tulp4* encodes an endogenous circRNA, we designed convergent and divergent primers that amplify canonical or back-spliced forms of *Tulp4* RNA (Fig. 2B, upper panel). PCR after reverse-transcription with the divergent primers (black) generated an RNase-R-resistant form of circTulp4, whereas the convergent primers (red) generated an RNase-R-sensitive product from linear *Tulp4* mRNA (Fig. 2B, lower panel). Moreover, Sanger sequencing confirmed the junction sequence in circTulp4 (Fig. 2C).

We next precisely positioned and quantified circTulp4 by using BaseScope (a novel mutation-specific RNA ISH assay); we used probes designed to target the circTulp4 junction sequence or a negative-control RNA sequence (bacterial *dap*B mRNA). Probe details are listed in Supplementary Data. BaseScope signals were readily distinguishable as punctate red dots within cells, and circTulp4 expression was markedly lower in 9-month-old APP/PS1 mice than WT mice (Fig. 2D); the difference was not notable in 6-month-old mice.

In FISH analysis performed for cellular localization, a junction-specific probe detected abundant nuclear circTulp4 expression in Neuro-2a cells (Fig. 2E); moreover, nuclear mass-separation assays revealed that in primary hippocampal neurons, circTulp4 was mostly localized in the nucleus (Fig. 2F). Because the levels of certain circRNAs (including intron lariats) are positively correlated with those of their parental mRNAs [23], we examined whether circTulp4 exhibits this relationship: siRNA-mediated knockdown of circTulp4 resulted in a reduction in *Tulp4* mRNA and protein levels in primary hippocampal neurons (Fig. 2G); the same effect was produced by three circTulp4-specific siRNAs (Supplementary Fig. 2A-B). Thus, when circTulp4 levels were lowered, *Tulp4* expression was diminished; however, siRNA-mediated knockdown of *Tulp4* mRNA did not affect circTulp4 levels (Supplementary Fig. 2C). Conversely, *Tulp4* mRNA and protein levels were increased when circTulp4 was overexpressed in cells (Fig. 2G). Collectively, these results suggest that circTulp4 can regulate *Tulp4* expression.

### CircTulp4 associates with RNA polymerase II by interacting with U1 snRNP

To investigate how circTulp4 regulates *Tulp4*, we examined circTulp4-protein interactions. First, we evaluated circTulp4-protein interaction tendency by using CatRAPID, which yields Interaction Strengths indicating likely occurrence (>50%) or high-confidence prediction (>75%) of interaction [21] (Section 2.14). Notably, an Interaction Strength of 99% was obtained for circTulp4 interaction with SNRPA (U1 snRNP) (Fig. 3A).

**Figure 3.**
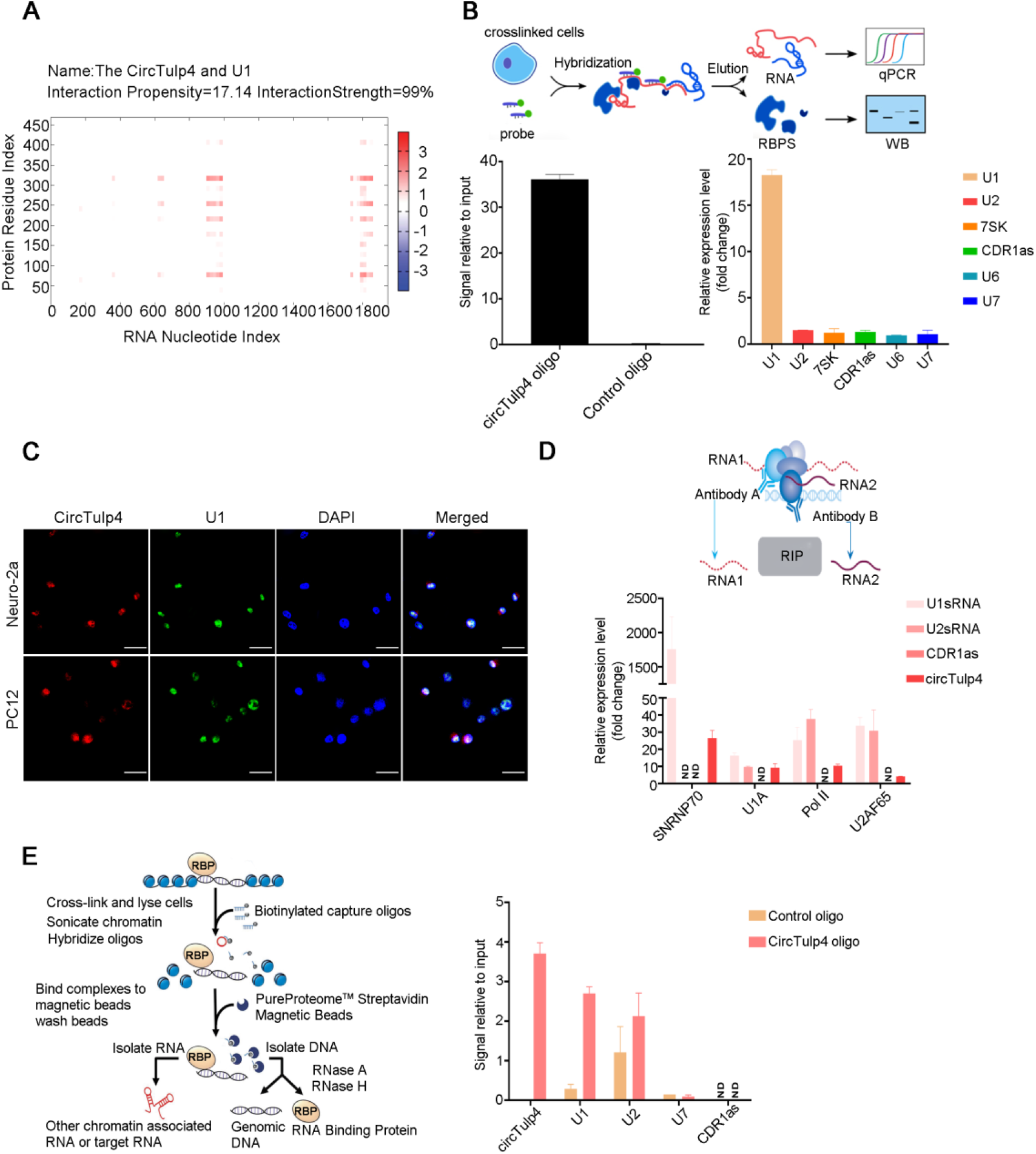
CircTulp4 associates with RNA polymerase II by directly interacting with U1 snRNP. **a** Heatmap showing prediction results for U1 and circTulp4 interaction; x-axis and y-axis: indexes of RNA and protein sequences, respectively. Heatmap colors indicate interaction scores, from −3 to 3, of individual amino acid and nucleotide pairs. The total sum represents the overall interaction score. The algorithm CatRAPID identified interaction between U1 and circTulp4 with confidence: Interaction Propensity, 17.14; Interaction Strength, 99%. **b** RNA antisense purification (RAP) assays were used to pulldown RNAs with circTulp4; U2 snRNA, CDR1as, U6 snRNA, U7 snRNA, and 7SK are controls. The circTulp4 probe captured considerably more circTulp4 relative to the control probe, and U1 was pulled down to a significantly higher level than other RNAs; this suggests that circTulp4 can bind to U1 snRNA. **c** Double FISH for circTulp4 and U1. Nuclei were stained with DAPI; circTulp4 was labeled with an Alexa Fluor 555-conjugated probe; U1 snRNA was labeled with an FITC-conjugated probe. Original magnification 40×; scale bar : 20 μm. **d** RNA immunoprecipitation (RIP) assay with antibodies against U1A, SNRNP70, and RNA polymerase II (Pol II). U2AF65 is shown for comparison. U1A, SNRNP70, and Pol II co-precipitated a substantial amount of circTulp4 but not CDR1as; pulldown with U2AF65 did not result in marked enrichment of circTulp4. ND: not detected. **e** Pulldown of indicated RNAs in chromatin isolation by RNA purification (ChIRP) assays for circTulp4. Pulldown with circTulp4 probes resulted in capture of U1 snRNA. U2 snRNA was also pulled down with circTulp4, although the association between U2 snRNA and circTulp4 could be indirect. Results are shown as means ± SD.

We next used RAP assays to directly test circTulp4 interaction with U1 snRNA (Fig. 3B). Because U1 regulatory transcription has been demonstrated [24], U1 snRNP was considered likely to bind to circTulp4, and thus we designed primers against U1. Because U1 and U2 are generally in close proximity and frequently function together to form a splicing body, we checked for U2 capture. We selected 7SK and CDR1as because 7SK has been reported to function in regulating transcription [25], and CDR1as has been comprehensively investigated in the nervous system and in degenerative diseases [26]. Lastly, U6 and U7, which are distant from U1, were used as negative controls. The circTulp4-specific probe captured considerably more circTulp4 relative to the control probe, and U1 was pulled down to a markedly higher level than other RNAs (Fig. 3B). Notably, U1 snRNA also colocalized with circTulp4 (detected using a junction probe) in two cell types (Fig. 3C). These results suggest that circTulp4 can bind to U1 snRNA.

Because U1 interacts with RNA polymerase II to regulate transcription [24], we tested whether circTulp4 interacts with the U1 protein complex and RNA polymerase II. In RIP assays performed using antibodies against U1 (U1A and SNRNP70), RNA polymerase II, and U2, RNA polymerase II showed almost no binding to CDR1as but associated with circTulp4 (Fig. 3D). This provides complementary evidence that these molecules interact and thus might function together.

We also investigated potential interactions between circTulp4, U1 snRNA, and *Tulp4*; we checked for RNA capture by performing ChIRP assays, which revealed that pull-down with the circTulp4 probe captured U1 snRNA (Fig. 3E). U2 snRNA was also pulled down with circTulp4, although this association might be indirect. These results verified the RAP results and demonstrated circTulp4 association with U1 snRNA.

### CircTulp4 recruits U1 snRNP to regulate Tulp4 transcription

We divided the 3000-bp *Tulp4* promotor (3000 bp upstream of coding area) into 6 segments (A–F) and designed a primer for almost every 500-bp stretch. We found that circTulp4 associates with segments E and F, located ~500 bp upstream of the transcriptional start site of *Tulp4* (Fig. 4A).

**Figure 4.**
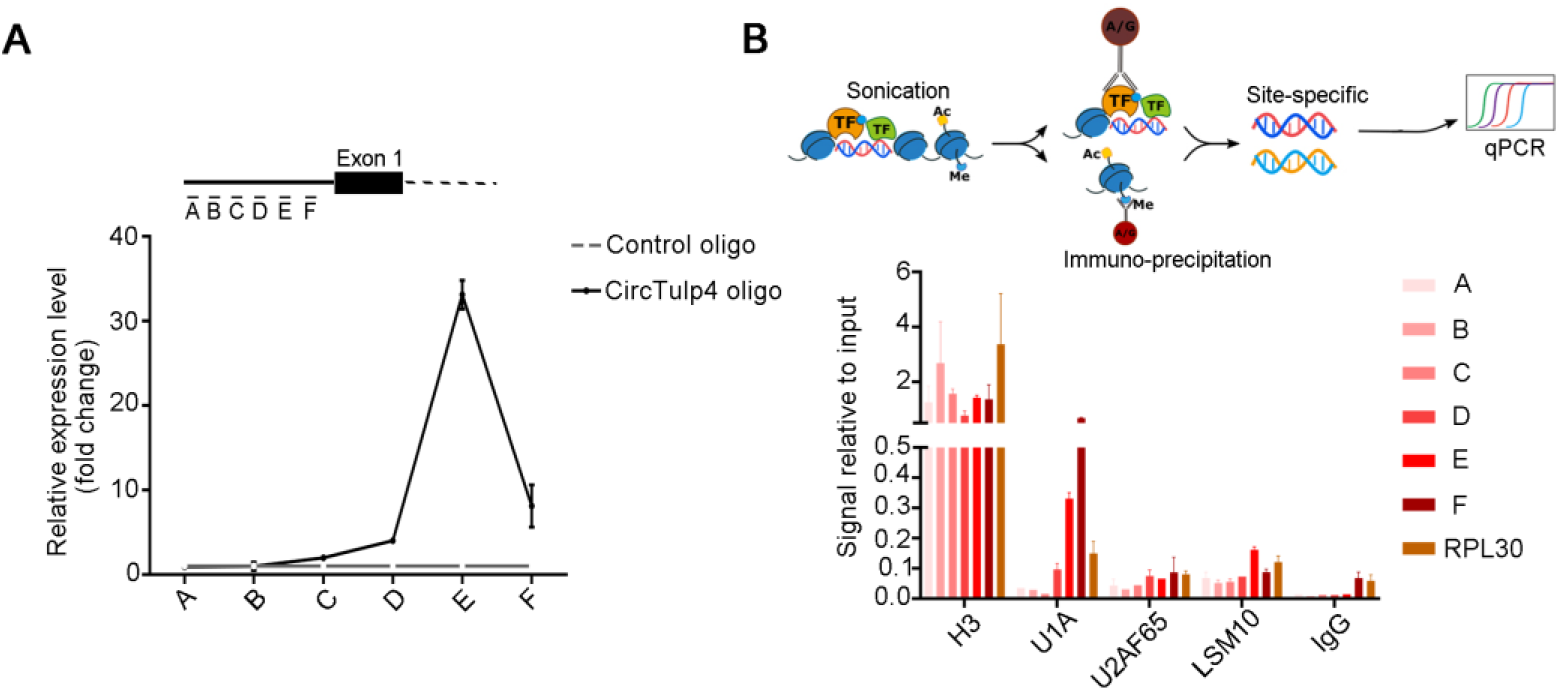
CircTulp4 recruits U1 snRNP and thereby regulates *Tulp4* transcription. **a** Pulldown of parental-gene promoter in ChIRP for circTulp4. The results indicate that circTulp4 occupies a region ~500 bp upstream of the transcriptional start site of *Tulp4*. **b** Pulldown of parental-gene promoter in ChIP assay performed using antibodies against U1A. For comparison, results are shown for U2AF65 and LSM10 (a protein component of U7 snRNP). IgG, negative control; H3, positive control. The results show that U1 snRNP binds to *Tulp4* promoter, indicating that circTulp4 and U1 snRNP (U1A) occupy a region ~500 bp upstream of *Tulp4* transcriptional start site; this suggests that circTulp4, U1 snRNP, and Pol II potentially interact with each other at the promoter region of the parental gene. Results are shown as means ± SD.

Next, we performed *Tulp4* promoter pull-downs in ChIP assays by using a U1A antibody, with the assays including antibodies against control molecules U2AF65 and LSM10 (a protein component of U7 snRNP) and positive- and negative-control antibodies (anti-H3 histone and IgG, respectively). RPL30 was the positive-control primer, and chromatin was ultrasonically broken into 200–500-bp fragments (Supplementary Fig. 3). Segments E and F showed marked enrichment, which indicated that the U1A-binding site was located in these segments, and U1A but not U2AF65 interacted with the *Tulp4* promoter region (Fig. 4B), which indicated that U1 snRNP specifically binds to *Tulp4* promoter. These findings suggest that circTulp4, U1 snRNP, and RNA polymerase II potentially interact with each other at the promoter region of *Tulp4*.

### CircTulp4 affects Tulp4 expression and thereby regulates neuronal differentiation, cell viability, and neurite growth and branching

To complete our current evaluation of circTulp4 as a potential AD biomarker, we examined circTulp4 neuronal expression and function. First, qPCR results demonstrated circTulp4 expression in brain tissue from 2–12-month-old APP/PS1 and WT mice, and in accord with the RNA-seq results, circTulp4 expression in the brain was lower in APP/PS1 than WT mice at both 9 and 12 months (Fig. 5A). Moreover, in agreement with a previous study [11], circTulp4 was upregulated during neuronal differentiation in different cell types (Fig. 5B, Supplementary Fig. 4A).

**Figure 5.**
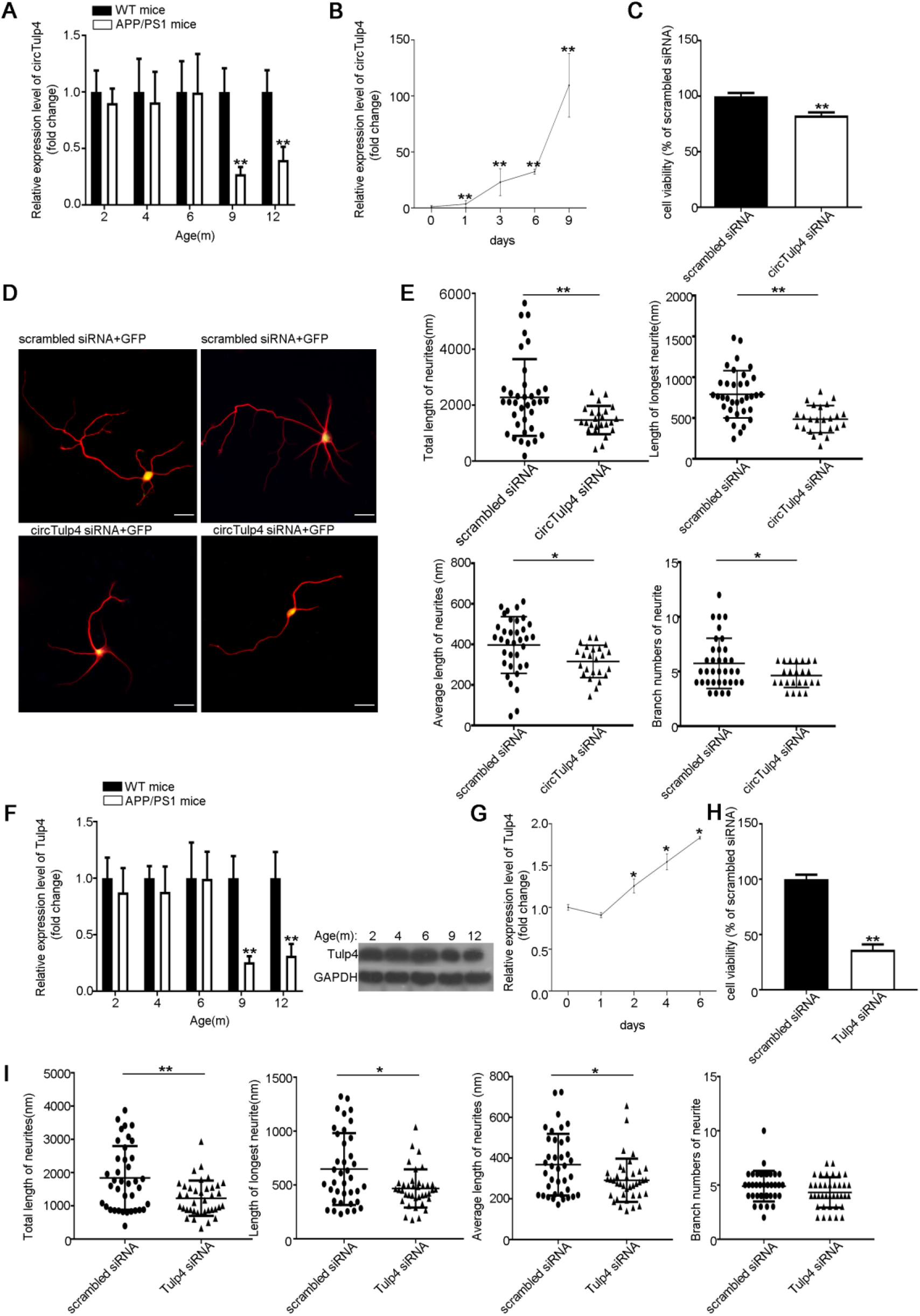
CircTulp4 regulates neuronal differentiation, cell viability, and neurite growth and branching through its parental gene, *Tulp4*. **a** Relative expression levels of circTulp4 in WT and APP/PS1 mouse cortexes were examined using qRT-PCR (9 pairs of mice at each stage). Glyceraldehyde-3-phosphate dehydrogenase gene (*Gapdh*) was used as the reference gene. **b** qRT-PCR analysis of relative expression of circTulp4 in primary hippocampal neurons during development (n=3). **c** Mouse primary hippocampal neurons were transfected with circTulp4 siRNA (100 nM), and 72 h later, CCK-8 assay was performed to evaluate cell viability. **d,e** Mouse primary hippocampal neurons were co-transfected with PT-GFP and circTulp4 siRNA (100 nM) (concentration ratio 1:5) before culturing, and 72 h later, neurites were immunostained with anti-β-tubulin III antibody. The length and branch numbers of GFP-expressing neurites were quantified using ImageJ software. Original magnification 40×; scale bar : 40 μm. **f** Relative expression of Tulp4 gene and protein in WT and APP/PS1 mouse cortex: qRT-PCR and western blotting analyses (9 pairs of mice at each stage); the results were normalized relative to GAPDH. **g** qRT-PCR analysis of *Tulp4* relative expression in primary hippocampal neurons during development (n=3). **h** Mouse primary hippocampal neurons were transfected with Tulp4 siRNA (100 nM), and 72 h later, CCK-8 assay was performed to evaluate cell viability. **i** Mouse primary hippocampal neurons were co-transfected with PT-GFP and Tulp4 siRNA (100 nM) (concentration ratio 1:5) before culturing, and after culturing for 72 h, neurites were immunostained with anti-β-tubulin III antibody. The length and branch numbers of GFP-expressing neurites were quantified using ImageJ software. Results are shown as means ± SD (*p<0.05, **p<0.01).

To investigate circTulp4 function in neuronal cells, circTulp4 expression was silenced using junction-sequence-targeting siRNAs; these siRNAs, but not a control siRNA, markedly lowered circTulp4 levels in two cell types (Supplementary Fig. 2) and potently suppressed cell proliferation (Fig. 5C, Supplementary Fig. 4B). Moreover, circTulp4 downregulation significantly decreased neurite growth and branching in hippocampal neurons and cell lines (Fig. 5D-E, Supplementary Fig. 4C). These findings suggest that circTulp4 functions in neuronal growth and development.

Notably, Tulp4 exhibited the same expression pattern as circTulp4 in mouse brain tissue (Fig. 5F) and was upregulated during neuronal differentiation (Fig. 5G). The finding that circTulp4 knockdown affected *Tulp4* expression suggested that *Tulp4* plays a role in the effects observed following circTulp4 knockdown. Accordingly, siRNA-mediated *Tulp4* downregulation markedly diminished neuronal viability (Fig. 5H, Supplementary Fig. 4D), and Tulp4 knockdown in primary hippocampal neurons also strongly inhibited neurite growth and branching (Fig. 5I). And the subsequent rescue experiment can be found that when knocking down circTulp4 and overexpressing the parent gene Tulp4 at the same time, it has a significant rescue effect on cell viability and neurite damage (Fig. 6A-B). These findings collectively suggest that circTulp4 could influence normal neuronal growth and development by regulating *Tulp4* expression; we propose that this circTulp4 regulatory function underlies the potential link between circTulp4 dysregulation and AD pathogenesis.

**Figure 6.**
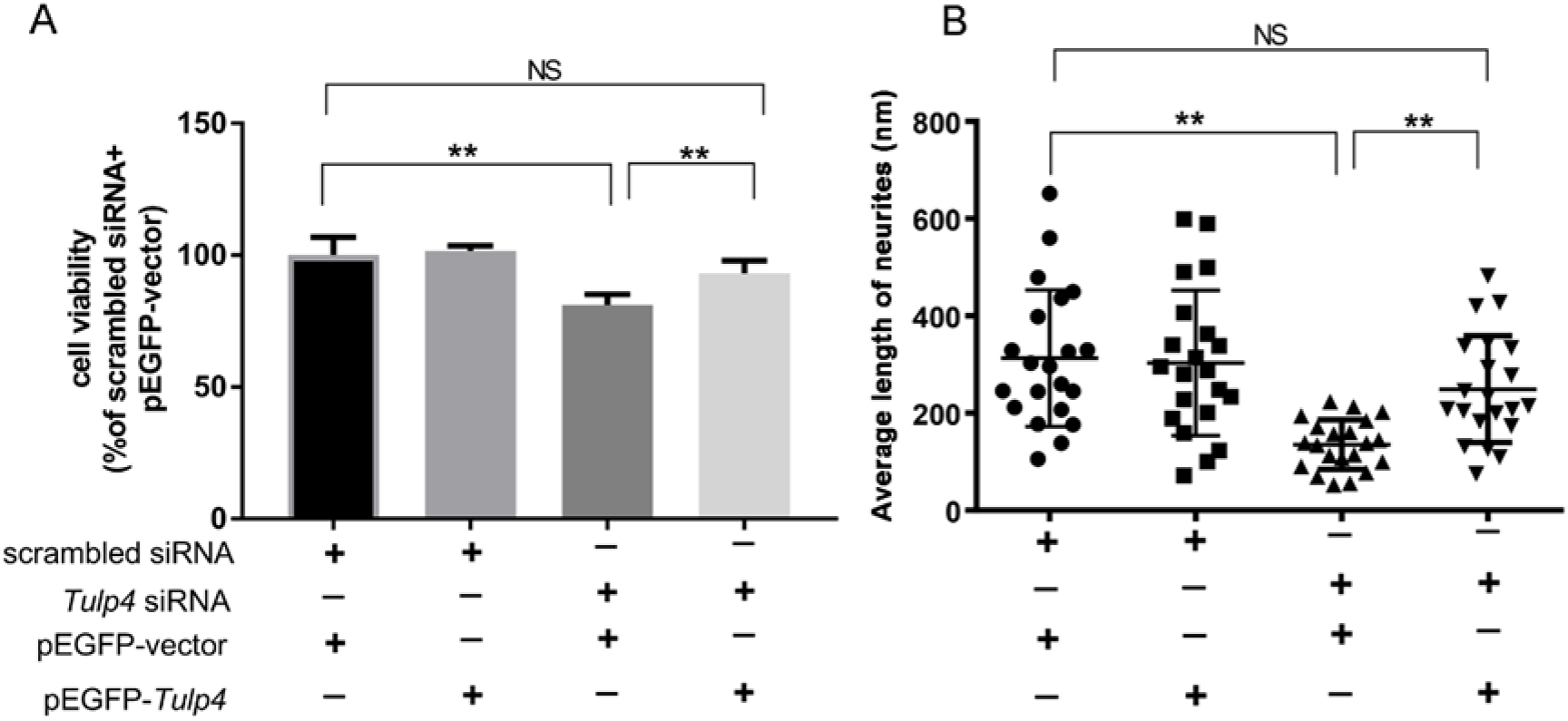
CircTulp4 regulates neuronal differentiation, cell viability, and neurite growth and branching through its parental gene, Tulp4 (rescue experiment) It can be found that when knocking down circTulp4 and over-expressing the parent gene Tulp4 at the same time, there is an obvious rescue effect. **a** Mouse primary hippocampal neurons were co-transfected with pEGFP-Tulp4 and circTulp4 siRNA before culturing, and after culturing for 72 h, CCK-8 assay was performed to evaluate cell viability. **b** Mouse primary hippocampal neurons were co-transfected with pEGFP-Tulp4 and circTulp4 siRNA before culturing, and after culturing for 72 h, neurites were immunostained with anti-β-tubulin III antibody. The length and branch numbers of GFP-expressing neurites were quantified using ImageJ software. Results are shown as means ± SD (*p<0.05, **p<0.01).

## Methods

### Animals

WT and APP/PS1 mice (The Jackson Laboratory; strain B6.Cg-Tg(APPswe, PSEN1dE9)85Dbo/Mmjax) were purchased from the Model Animal Research Center of Nanjing University, housed 1/cage under standard specific conditions (25°C, 50% humidity, 12/12-h light/dark cycle, pathogen-free environment), and provided free access to food until 6/9 months old. From 3 randomly selected male mice from each group, brain tissues were collected for RNA-seq. All animal experiments were performed in accordance with animal use protocols approved by the Committee for the Ethics of Animal Experiments, Shenzhen Peking University The Hong Kong University of Science and Technology Medical Center (SPHMC) (protocol number 2011-004).

#### RNA-seq

Total RNA was isolated by using TRIzol reagent (Invitrogen) as per manufacturer instructions. After RNA isolation, RNA degradation and contamination were assessed using (1%) agarose-gel electrophoresis, RNA purity was checked using a NanoPhotometer spectrophotometer (IMPLEN, CA, USA), RNA concentration was measured using a Qubit RNA Assay Kit and Qubit 2.0 Fluorometer (Life Technologies, CA, USA), and RNA integrity was evaluated using the RNA Nano 6000 Assay Kit for a Bioanalyzer 2100 System (Agilent Technologies, CA, USA). The complete circRNA study involved the following steps: RNA isolation, quantification, and qualification; library preparation for circRNA sequencing; sequencing and clustering of reads; quality control; mapping to the reference genome; circRNA identification; quantification of gene-expression level; and differential-expression analysis. Normalized circRNA levels (estimated based on TPM values) were calculated as listed below, and transcripts were considered to be differentially expressed between APP/PS1 and WT mice when their p value was <0.05.

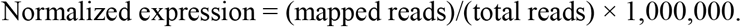

#### Cell culture

Neurons were isolated from the brain of E16.5–E18.5 mice (C57B6), seeded on petri dishes coated with poly-D-lysine (Sigma), and cultured in neurobasal medium (Life Technologies) containing 2% B27 (Life Technologies) and 0.5 mM L-glutamine at 37°C in a 5% CO2 incubator. The cell lines Neuro-2a (mouse neuroblastoma N2a cells) and HEK293T were grown in DMEM (GIBCO) supplemented with 10% FBS (HyClone) and 1% antibiotics, and PC12 cells (rat pheochromocytoma cells) were grown in DMEM supplemented with 6% FBS, 6% HS (HyClone), and 1% antibiotics. Differentiation was induced (for 48 h) by maintaining Neuro-2a cells in DMEM containing 0.5% FBS, 10 mM RA (retinoic acid; Sigma), and 1% antibiotics, and PC12 cells in DMEM containing 0.5% FBS, 0.5% HS, 100 ng/mL NGF (nerve growth factor; Sigma), and 1% antibiotics.

#### Transfection and RNA interference

Neuro-2a cells seeded in 6-well plates (6×10^5^/well) were transfected with 100 nM circTulp4-specific or negative-control siRNA by using Lipofectamine® RNAiMAX (Invitrogen). At 24 h post-transfection, cells were lysed, RNA was isolated, and siRNA transfection efficiency was tested using quantitative real-time RT-PCR (qRT-PCR). Differentiation was induced for 48 h. Dissected hippocampal neurons seeded in 12- or 24-well plates (1×10^5^/well or 5×10^4^/well) were co-transfected with 50 or 25 nM circTulp4 siRNA and EGFP plasmid (5:1 molar ratio); and immunofluorescence analysis was performed at 72 h post-transfection.

#### qRT-PCR

Total RNA was extracted using TRIzol reagent (Sigma), according to the manufacturer’s protocol, and RNA quantity was measured using a NanoDrop 2000 (Thermo Fisher Scientific). qRT-PCR was performed using the GoScriptTM Reverse Transcription System (Promega) and a C1000 Thermal Cycler (Bio-Rad). Glyceraldehyde-3-phosphate dehydrogenase gene (Gapdh) was used as an internal control. Relative gene-expression levels were quantified using the 2−ΔCt method.

#### Cell counting kit-8 (CCK-8) assay

Treated cells were seeded in 24-well plates (1×10^4^/well) and incubated in a 5% CO_2_ incubator at 37°C, and then CCK-8 solution (Cat. no. HY-K0301-100T; MedChem Express, Monmouth Junction, NJ, USA) was added (50 μL/well). After 3/4-h incubation, optical density was measured using a microplate reader (Bio-Rad). Cell viability is presented as a percentage relative to nontreated cells.

#### Fluorescence in situ hybridization (FISH)

Cy3-labeled circTulp4 probes were designed and synthesized by RiboBio (Guangzhou, China; probe sequences available upon request), and probe signals were detected using a FISH kit (RiboBio), as per manufacturer instructions. Cells were stained with 4′-6-diamidino-2-phenylindole (DAPI; Life Technologies, Carlsbad, CA, USA) before examination under a fluorescence microscope (Zeiss LSM 710).

#### Immunofluorescence analysis

Neurons were fixed in 4% (w/v) paraformaldehyde (30 min, room temperature) and then incubated with PBS containing 0.4% Triton X-100, 1% bovine serum albumin (Sigma), and 4% goat serum for 30 min to permeabilize the cells and block nonspecific staining. Neurons were incubated overnight at 4°C with a tubulin primary antibody (mouse monoclonal anti-β-tubulin III, 1:500, Sigma) and for 2 h with a CY3-conjugated anti-mouse secondary antibody (1:500, Jackson Immuno Research), and then the coverslips were mounted on slides with an antifade mounting medium (Beyotime) and examined under a fluorescence microscope.

#### Western blotting

Protein extracts were prepared in RIPA buffer in the presence of a protease-inhibitor cocktail (Sangon) and 1 mM phenylmethanesulfonylfluoride (Sigma). After determining protein concentrations by using the Bradford protein assay, samples were separated using 10% SDS-PAGE and transferred to nitrocellulose membranes, which were probed with primary antibodies (4°C, overnight) and then with horseradish peroxidase-conjugated secondary antibodies (2 h). Signals were developed using a Western Lightning PLUS kit (NEL105001EA, PerkinElmer) and quantified using Quantity One (Bio-Rad).

#### In situ hybridization (ISH)

We used the single-pair probe ISH approach (BaseScope, Advanced Cell Diagnostics, Newark, CA, USA) based on the multiplex fluorescent ISH RNAscope® method (Advanced Cell Diagnostics [47]). ISH methods are highly specific and sensitive due to the use of a unique probe design employing “ZZ” probe pairs and signal amplification, respectively. Advances in signal amplification over RNAscope® have enabled the use in BaseScope assays of a single-pair probe, comprising a pair of 18–25-bp oligonucleotides, for detecting junction sequences. Table S1 lists the targeted sequences of our customized junction-specific circTulp4 ISH probes. BaseScope ISH assays were performed on 20-μm-thick, fresh-frozen brain-tissue sections from 6-/9-month-old WT and APP/PS1 mice. Sections were deparaffinized in xylene, treated with H_2_O_2_ (10 min, room temperature) to block endogenous peroxidases, permeabilized by performing antigen retrieval (15 min, 100°C), and incubated with a protease mixture (30 min, 40°C). Probes were bound through incubation for 2 h at 40°C, chemically amplified, and labeled using fluorophores (multiplex ISH) or through alkaline-phosphatase conversion of Fast RED dye (single-pair probe ISH). Table S1 lists the probe sequences.

#### RNA antisense purification (RAP)

RAP kits (Bersin Bio) were used according to manufacturer instructions. RAP employs specific biotinylated probes that hybridize to target RNAs, which can be pulled down, reverse-transcribed into cDNA, and identified through qRT-PCR analysis or sequencing. Here, 107 cells were washed with PBS, crosslinked using formaldehyde, lysed with 1 mL of lysis buffer, and homogenized using a 0.4-mm syringe, after which a single 50-bp biotinylated antisense probe (0.2 nmol) targeting the adaptor sequence was added. Probes were denatured (65°C, 10 min) and hybridized (room temperature, 2 h), and then 200 μL of streptavidin-coated magnetic beads were added. After washing to remove nonspecifically bound RNAs, TRIzol reagent was used to recover RNAs that specifically interacted with circTulp4. Table S1 lists the probe sequences.

#### RNA immunoprecipitation (RIP)

RIP assays were performed using a kit from Millipore (Cat. no. 17-701) and following the manufacturer’s instructions. Neuro-2a cells (2×10^7^) were lysed in complete RIP lysis buffer and the cell lysates were divided into equal parts and incubated (overnight with rotation, at 4°C) with anti-Pol II (Cell Signaling Technology (CST), Cat. no. #2629), anti-U1A (Abcam, Cat. no. ab166890), anti-U2AF65 (Abcam, Cat. no. ab37530), IgG (Millipore, Cat. no. pp64B), or anti-SNRNP70 (Millipore, Cat. no. CS203216). Next, magnetic beads (Millipore, Cat. no. CS203178) were added to the cell lysates and the incubation was continued at 4°C for 1 h, after which the samples were incubated with proteinase K at 55°C for 1 h. The enriched RNA was obtained using RNA Extraction Reagent (phenol:chloroform:isoamylol = 125:24:1; pH<5.0; Solar Bio), and qRT-PCR was performed on the purified RNA to detect the targets of interest.

#### Chromatin isolation by RNA purification (ChIRP)

ChIRP was performed using a kit from Millipore (Cat. no. 17-10495), according to the manufacturer’s instructions. Briefly, Neuro-2a cells grown in differentiation medium for 24 h were trypsinized (0.25% trypsin) and washed with 1× PBS, and then 2×107 cells were crosslinked with 20 mL of 1% glutaraldehyde/PBS at room temperature (18°C –25°C) for 10 min on an end-to-end rotator. The excess glutaraldehyde was quenched by adding 2 mL of 1.25 M glycine and incubating for an additional 5 min. After washing with 20 mL of cold PBS, the crosslinked cells were resuspended in 2 mL of lysis buffer and sonicated (in a cold room) by using a Bioruptor (Diagenode) with the following parameters: H—high setting; pulse interval—30 s ON and 30 s OFF; 10 repeats. After sonication, the fragmented chromatin samples (200–500-bp long) were split into two parts and hybridized with BiotinTEG-labeled tiling probes against circTulp4 (Table S1) or control probes in hybridization buffer at 37°C; after hybridization for 4 h, 100 μL of Pure Proteome Streptavidin magnetic beads (Millipore, Cat. no. CS219080) were added into each reaction and incubated at 37°C for 30 min, and the beads were then washed 4 times with prewarmed washing buffer at 37°C for 5 min. The retrieved bead –RNA–protein–DNA complex was split into two parts: 1/10 for RNA isolation, 9/10 for DNA purification. Lastly, the retrieved RNA and DNA were used as templates for real-time quantitative PCR (qPCR), and the obtained data are presented as the percentages of input RNA and DNA, respectively.

#### Rapid prediction of RNA-protein interaction

CircTulp4-protein interactions were predicted using CatRAPID, an online algorithm that rapidly predicts RNA-protein interactions based on secondary structure, hydrogen bonding, and molecular interatomic forces. The obtained interaction propensity is a measure of the interaction probability between 1 protein (or region) and 1 RNA (or region). This measure is based on the observed tendency of ribonucleoprotein-complex components to exhibit specific physicochemical properties that can be used to make a prediction. Interaction Strength, a statistical measure for evaluating interaction propensity with respect to CatRAPID training, reflects prediction confidence, ranging from 0% (unpredictability) to 100% (predictability): values of >50% indicate that an interaction is likely to occur, values of >75% represent high-confidence predictions [21].

#### Chromatin immunoprecipitation (ChIP)

ChIP experiments were performed using a kit from CST (Cat. no. #56383) and following the manufacturer’s instructions. Briefly, cells were fixed on plates with 1% formaldehyde for 10 min at room temperature, and after quenching the crosslinking by adding 0.125 M glycine (5 min), the cells were pelleted at 800 × g. Cell pellets were resuspended in 1 mL of SDS lysis buffer (1% (w/v) SDS, 10 mM EDTA, 50 mM Tris-HCl, pH 8.1) containing Roche complete protease-inhibitor cocktail and incubated for 10 min on ice. Cell extracts were sonicated for 15 min with a Bioruptor (Diagenode) to obtain 500-bp DNA fragments. After saving a 100-μL sample of the supernatant as input, the remaining supernatant was diluted 1:10 in ChIP dilution buffer (0.01% (w/v) SDS, 1.1% (v/v) Triton X-100, 1.2 mM EDTA, 16.7 mM Tris-HCl, pH 8.1, 167 mM NaCl) containing protease inhibitors. The chromatin solution was precleared and then immunoprecipitated with antibodies against Pol II (CST, Cat. no. #2629), U1A (Abcam, Cat. no. ab166890), U2AF65 (Abcam, Cat. no. ab37530), LSM10 (Abcam, Cat. no. ab180128), or H3 (CST, Cat. no. #4620), or with (as an antibody control) IgG (CST, Cat. no. #2729). The immune complexes were eluted in 1% (w/v) SDS and 50 mM NaHCO3, and then the crosslinks were reversed for 6 h at 65°C. Samples were digested with proteinase K for 1 h at 45°C, and DNA was extracted using phenol/chloroform/isoamyl alcohol. The eluted DNA was subject to real-time qPCR to detect enriched genomic DNA regions.

#### Statistical analyses

Distribution normality was assessed using Kolmogorov–Smirnov test, implemented in PRISM statistical software (GraphPad). Multiple normally distributed groups were compared using ANOVA with Tukey post-test, two normally distributed groups were compared using *t* tests, and two non-normally distributed groups were compared using Mann–Whitney U test. High-throughput sequencing-related data were calculated and statistically computed using R software.

### Discussion

No effective AD treatment is currently available, and therapies targeting the main AD neuropathological hallmarks (β-amyloid aggregates, neurofibrillary tangles) have shown little success [27]. Thus, epigenetic regulation of AD pathogenesis has recently been widely studied to identify potential new biomarkers/therapeutic targets. Intriguingly, dynamic changes in DNA methylation and long noncoding RNAs in the brain are reported to contribute to AD [28,29]. However, potential circRNA functions in AD development/progression remain largely unknown.

RNA-seq analyses [3,30] have allowed detailed characterization of circRNAs, which present characteristics that could link them to neurodegenerative diseases. Thousands of circRNAs are highly expressed in the mammalian brain, accumulated in neural tissues during aging, conserved between rodents and humans, and more abundant than linear RNAs in neuronal synaptic fractions [11,31]. Thus, circRNAs could perform critical functions in the nervous system, and circRNA dysregulation might underlie neurodegenerative disease development/progression. Accordingly, Lukiw [14] and Zhao et al. [32] reported roles for circRNAs in AD. However, our study represents the first comprehensive circRNA profiling in the brain in the APP/PS1 mouse model of AD. We compared brain circRNA profiles between age-matched APP/PS1 and WT mice and identified several circRNAs showing altered expression in APP/PS1 mice. Moreover, we found marked circTulp4 downregulation in the brain in 9-/12-month-old APP/PS1 mice, which suggests a potential association of circTulp4 dysregulation with AD pathogenesis.

CircRNA functions have been extensively studied but remain incompletely understood. Certain circRNAs could function as miRNA sponges, but only a few circRNAs contain multiple binding sites for trapping specific miRNAs [10,33,34], and circRNAs might not necessarily act as miRNA sponges in human and mouse cells [11]. Interestingly, circRNA knockdown can cause diminished parental-gene expression, which supports the possibility that certain circRNAs upregulate RNA polymerase II-mediated transcription of parental genes [35]. Our findings suggest that circTulp4 is one such circRNA: circTulp4 predominantly localizes in the nucleus and interacts with U1 snRNP and RNA polymerase II to promote *Tulp4* transcription. This ability to control *Tulp4* expression might allow circTulp4 to regulate nervous system functions and thereby influence AD development.

We propose (Fig. 7) that circTulp4 and U1 snRNA bind through RNA-RNA interaction, and that the circTulp4-U1 snRNP complex interacts with the RNA polymerase II transcription complex at *Tulp4* promoter to enhance gene expression; once transcription is initiated, circTulp4 generation would further promote the gene’s transcription and create a positive-feedback loop.

**Figure 7.**
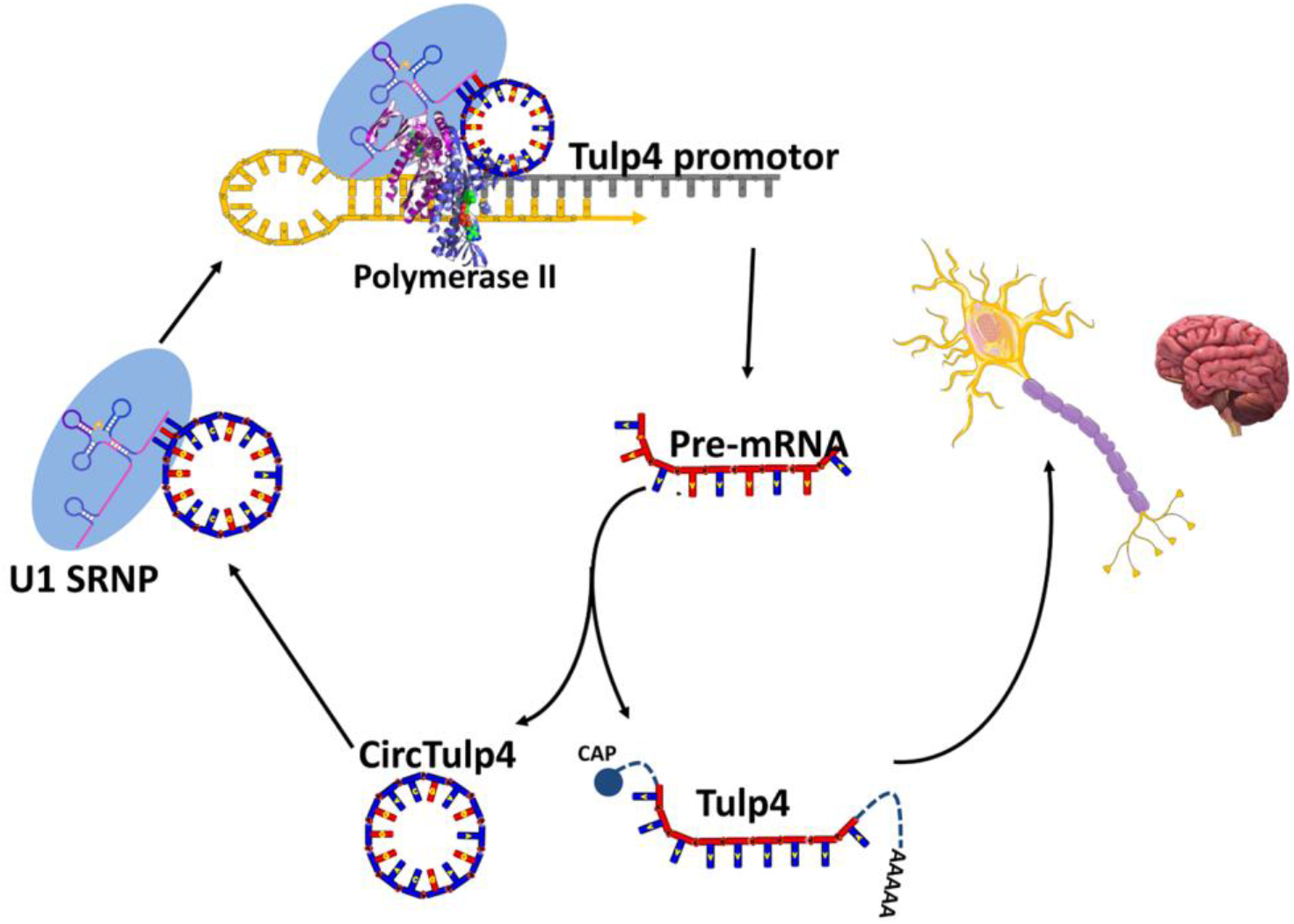
Working model depicting circTulp4 effects on *Tulp4* expression. CircTulp4 might form a complex with U1 snRNA through specific RNA-RNA interaction, and the circTulp4–U1 snRNP complex might further interact with the Pol II transcription complex at *Tulp4* promoter to modulate *Tulp4* expression and, thereby, regulate neuronal differentiation and nervous system functions and participate in AD development.

Tulp4 is a protein of the tubby family, which is associated with neuronal differentiation and development, and tubby-protein mutations are implicated in obesity, deafness, and visual impairment [36]: Mice carrying a mutated tubby gene develop delayed-onset obesity, sensorineural hearing loss, and retinal degeneration, and *TULP1* mutations cause retinitis pigmentosa in humans [37,38] and retinal degeneration in mice [39]. Moreover, Tulp3 mutants show embryonic lethality, with defects in dorsoventral patterning of the spinal cord [40,41]. However, very few studies have been published on Tulp4. This is the first report that Tulp4 can critically affect nerve-cell differentiation and thus potentially contribute to nervous system growth and development.

In conclusion, several brain-enriched circRNAs are differently expressed between WT and APP/PS1 mice; our RNA-seq database of these circRNAs could serve as a valuable resource for future investigations into circRNA involvement in AD development and suitability as AD pathogenesis biomarkers. Moreover, circTulp4 is downregulated in APP/PS1 mice, and circTulp4 localizes in the nucleus, interacts with U1 snRNP and RNA polymerase II, and promotes *Tulp4* transcription. The circTulp4 ability to control *Tulp4* expression might enable circTulp4 to regulate neuronal differentiation and thereby influence nervous system functions and AD development. Certain circRNAs are stably detected in body fluids (e.g., plasma), which allows their use as biomarkers or therapeutic targets [42–44]. Therefore, we will next test for stable circTulp4 detection in body fluids and determine whether circTulp4 expression correlates with specific clinical stages of AD.

### Conclusion

In conclusion, CircTulp4—a circRNA downregulated in APP/PS1 mice—localized in the nucleus and interacted with U1 snRNP and RNA polymerase II to regulate the transcription of its parental gene, Tulp4. CircTulp4 and Tulp4 showed similar expression in brain tissues, and downregulation suppressed neuronal survival/growth and branching. circTulp4 maight influence nervous system functions and might participate in the development of AD.

---

**Supplementary Fig 1.**
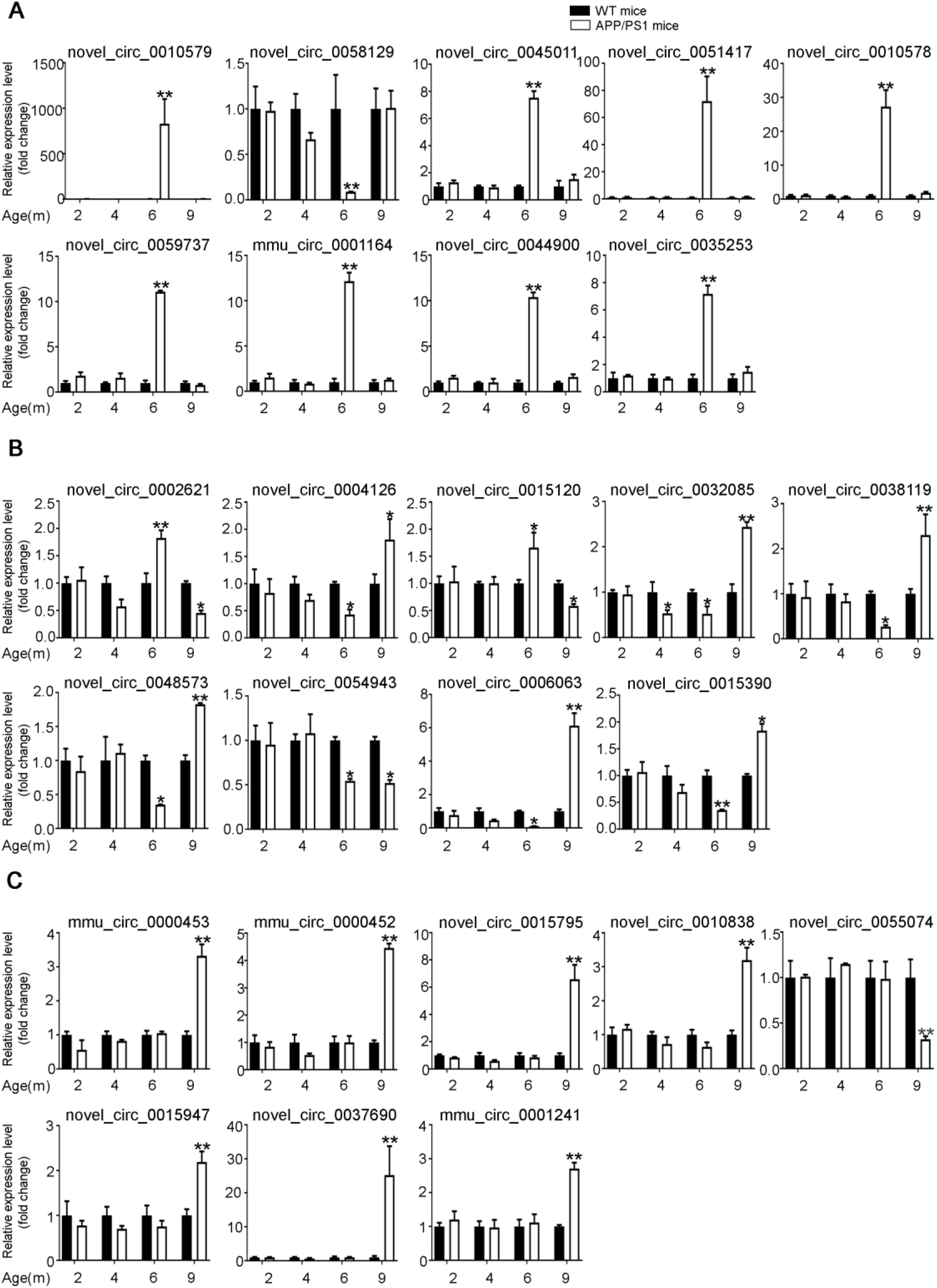
Quantitative PCR validation of circRNA expression. We selected three sets of circRNAs for verification: 6yes9no: differential expression at 6 months, no differential expression at 9 months; 6no9yes: no differential expression at 6 months, differential expression at 9 months; and 6yes9yes: differential expression at both 6 and 9 months. **a** 6yes9no group, **b** 6yes9yes group, **c** 6no9yes group. CircRNA expression was quantified relative to GAPDH expression by using the comparative cycle threshold (ΔCT) method. Data are presented as means ± SD (n=3, *p<0.05, **p<0.01).

**Supplementary Fig 2.**
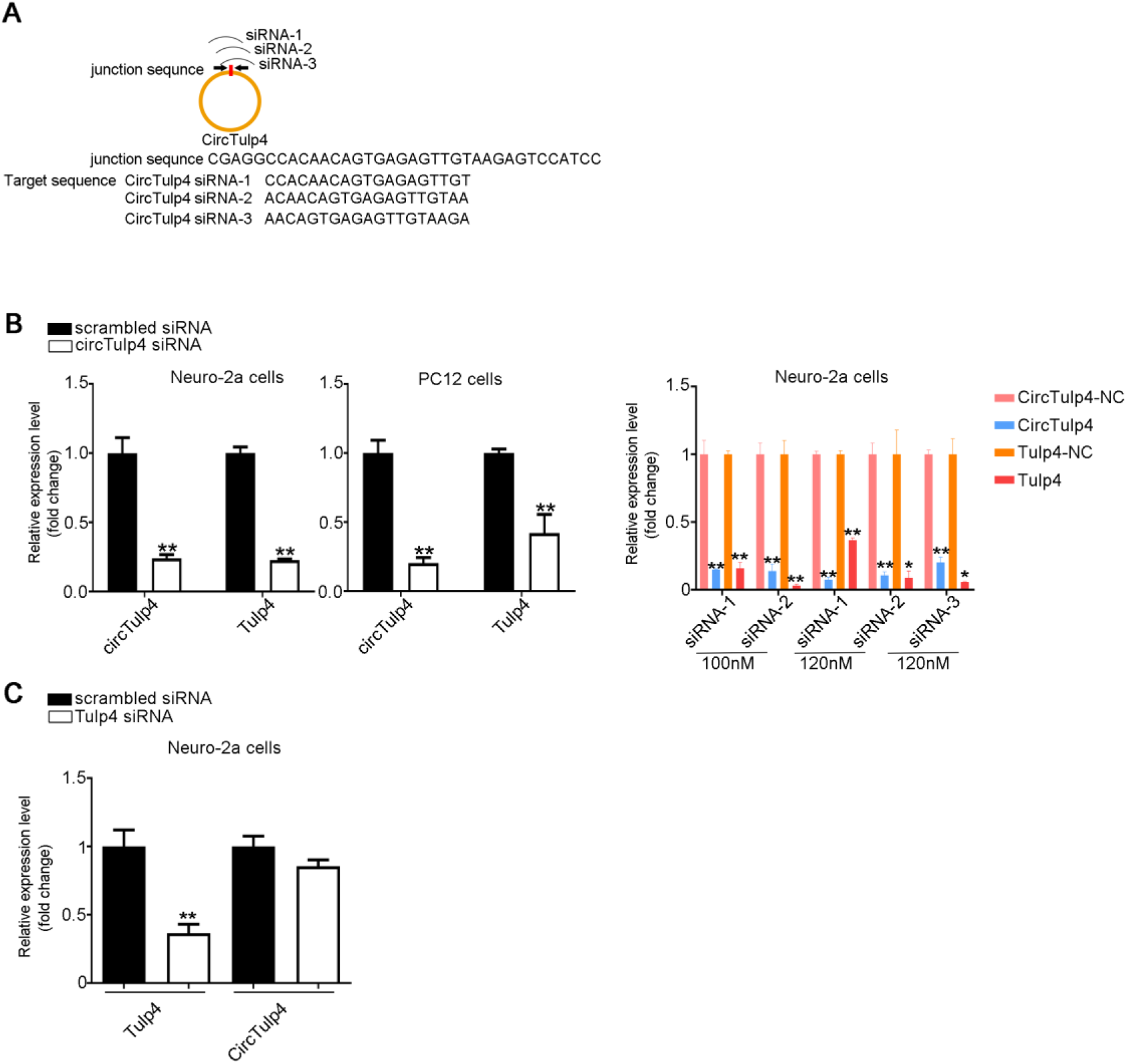
CircTulp4 is mainly localized in the nucleus and regulates its parental gene, *Tulp4*. **a** Target sequence for siRNAs. **b** Neuro-2a and PC12 cells were transfected with circTulp4 siRNA (100–120 nM) before culturing, and the cells were then cultured for 24 h. The qRT-PCR data indicate that the expression of the parental gene was decreased when circTulp4 was downregulated. **c** Neuro-2a cells were transfected with *Tulp4* siRNA (100 nM) before culturing, and the cells were then cultured for 24 h. The qRT-PCR data indicate that siRNA-mediated knockdown of *Tulp4* mRNA did not affect circTulp4 levels. Results are shown as means ± SD (*p<0.05, **p<0.01).

**Supplementary Fig 3.**
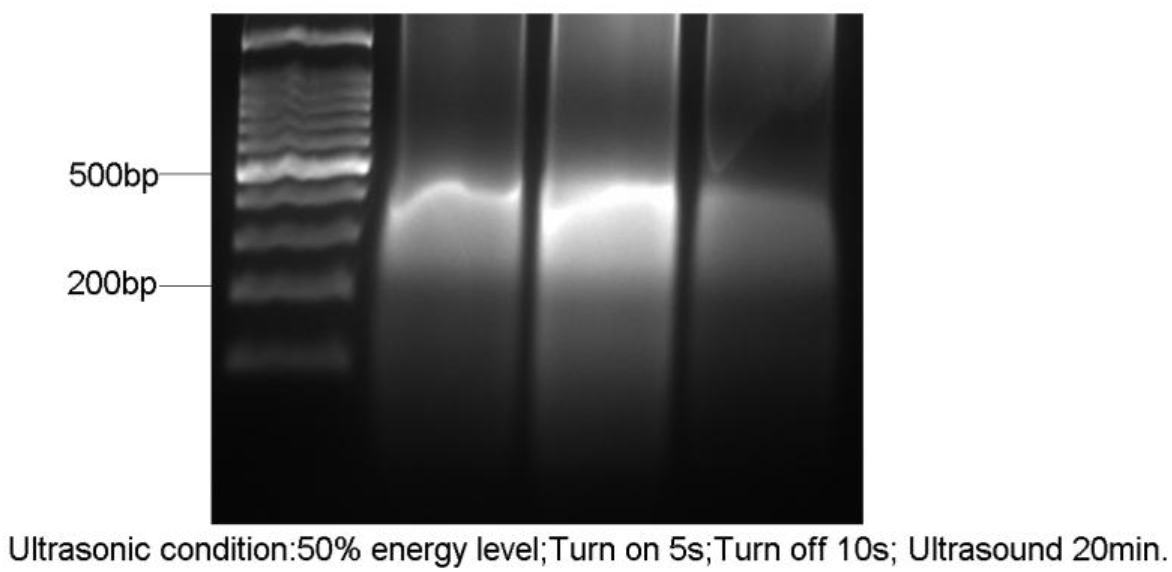
Chromatin fragmentation for ChIP analyses. Ultrasound conditions for breaking chromatin into 200–500-bp fragments: 50% energy level; turn on 5 s, turn off 10 s; ultrasound 20 min.

**Supplementary Fig 4.**
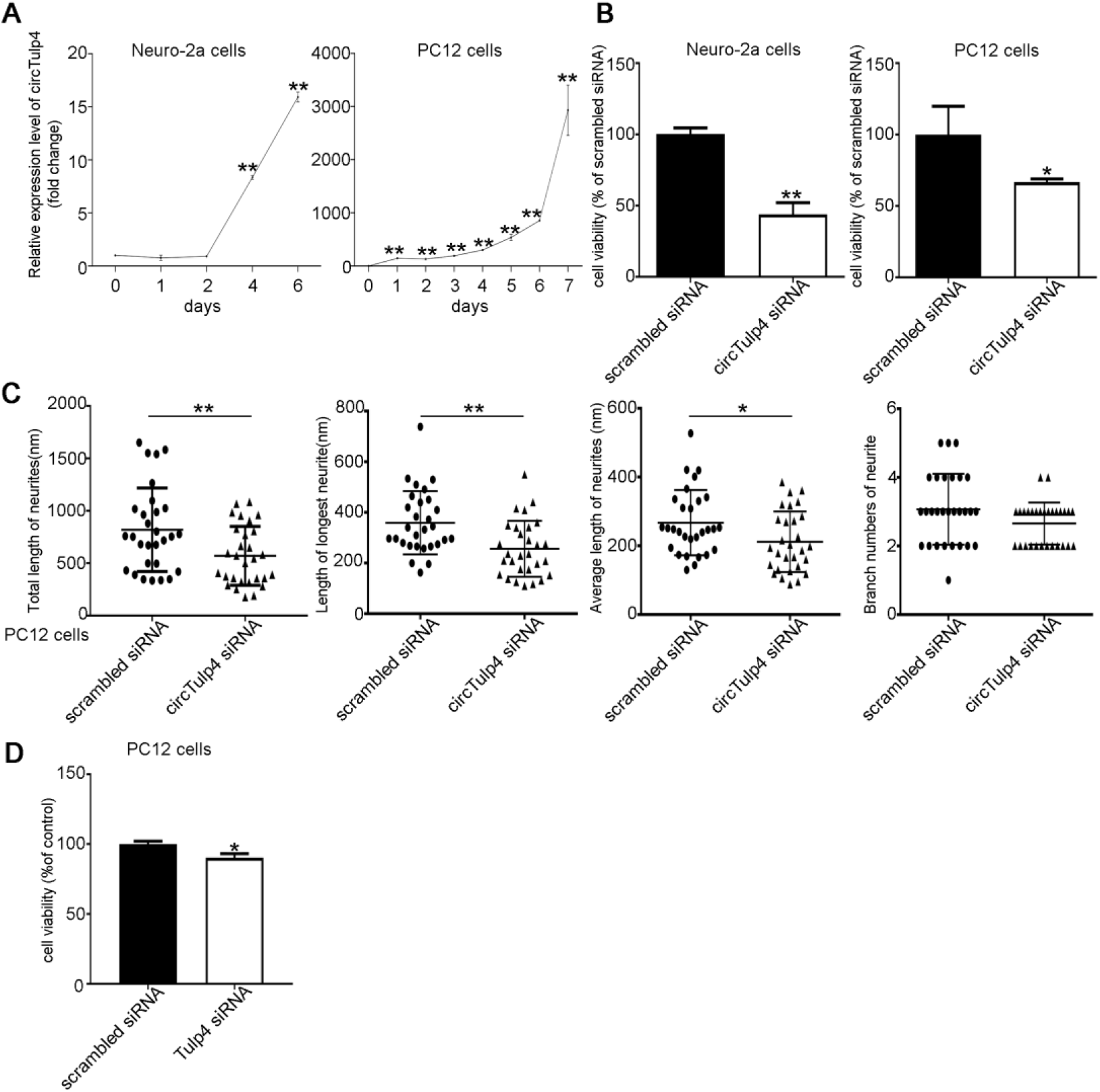
CircTulp4 regulates neuronal differentiation, cell viability, and neurite growth and branching through *Tulp4*. **a** qRT-PCR analysis of circTulp4 relative expression in Neuro-2a and PC12 cells during development in culture. **b** CCK-8 assay was performed to evaluate cell viability. Neuro-2a and PC12 cells were transfected with circTulp4 siRNA (100 nM) and then induced with RA (3 μg/mL) or NGF (50 μg/mL) for 48 h before use in CCK-8 assays. **c** PC12 cells were co-transfected with PT-GFP and circTulp4 siRNA (100 nM) (concentration ratio 1:5) before culturing, and after culturing for 48 h, the cells were treated with NGF (50 μg/mL) for 48 h. Neurites were immunostained with anti-β-tubulin III antibody. The length and branch numbers of GFP-expressing neurites were quantified using ImageJ software. **d** CCK-8 assay was used to evaluate cell viability. PC12 cells were transfected with Tulp4 siRNA (100 nM) and then treated with NGF (50 μg/mL); after 48-h induction, the cells were used in CCK-8 assays. Results are shown as means ± SD (*p<0.05, **p<0.01).

## Acknowledgments

We thank the Shenzhen Biomedical Research Support Platform and the Shenzhen Molecular Diagnostic Platform of Dermatology for technical help.

This work was supported by National Key R&D Program of China Grant 2016YFA0501903, National Natural Scientific Foundation of China (Grant Nos. 81673053, 81701069), Natural Scientific Foundation of Guangdong Province (2016A030312016), and Shenzhen Basic Research Grants (JCYJ20170411090739316, JCYJ20170815153617033, JCYJ20180507182657867, JCYJ20170306161713757).

## Data availability

Gene-expression data are available at the Gene Expression Omnibus (http://www.ncbi.nlm.nih.gov/geo/; Accession Number GSE-132177). All other data are available on request.

## Ethics approval and consent to participate

All animal experiments were performed in accordance with animal use protocols approved by the Committee for the Ethics of Animal Experiments, Shenzhen Peking University The Hong Kong University of Science and Technology Medical Center (SPHMC) (protocol number 2011-004).

## Author contributions

JW, WZ, and NM designed the study. NM, JP, QW and BY participated in the animal experiments including tissue collection and RNA/protein extraction. NM performed the experiments and collected the data. NM, BY, JW, and WZ analyzed the results and wrote the manuscript. All authors read and approved the final manuscript.

---

**Table S1.**
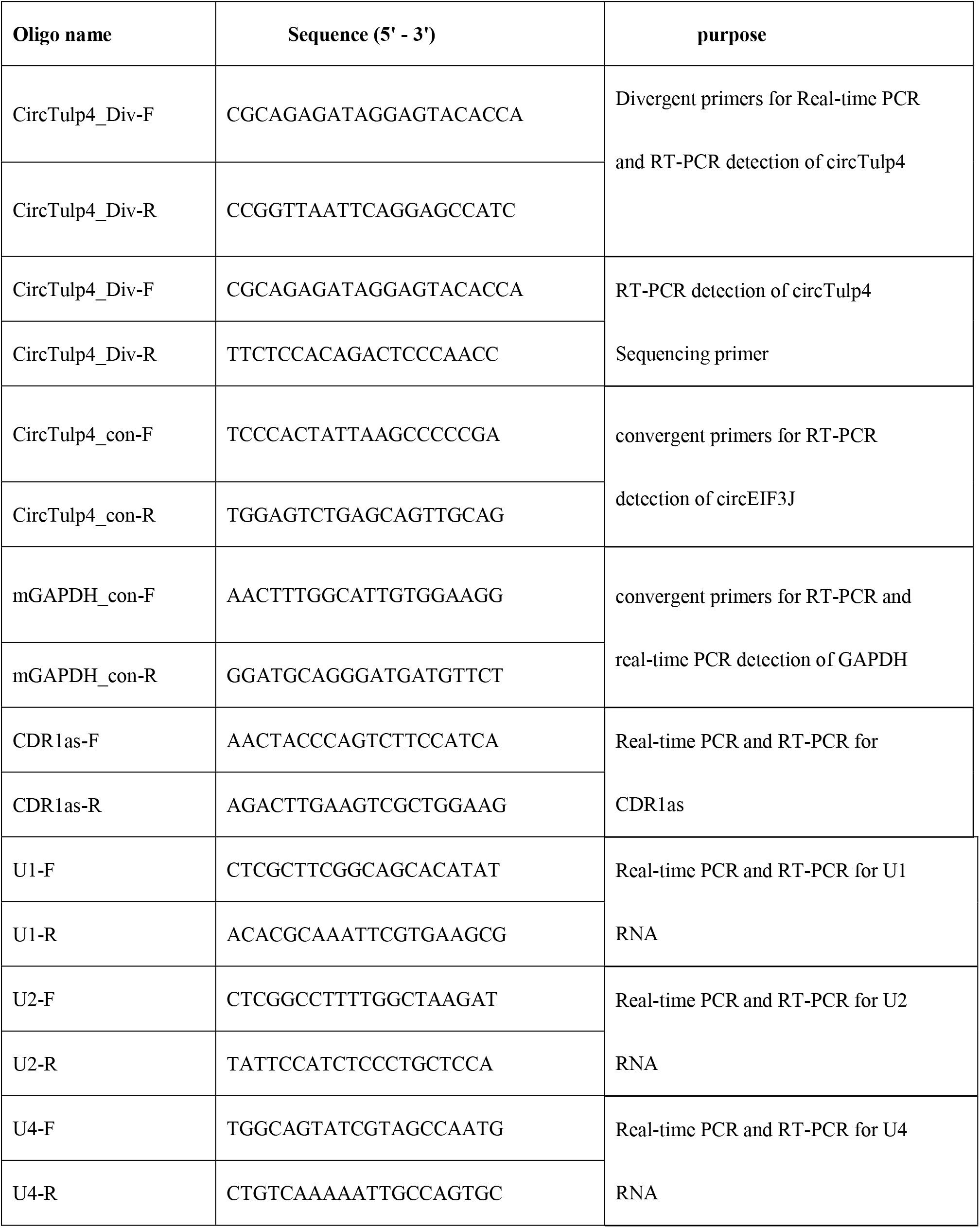
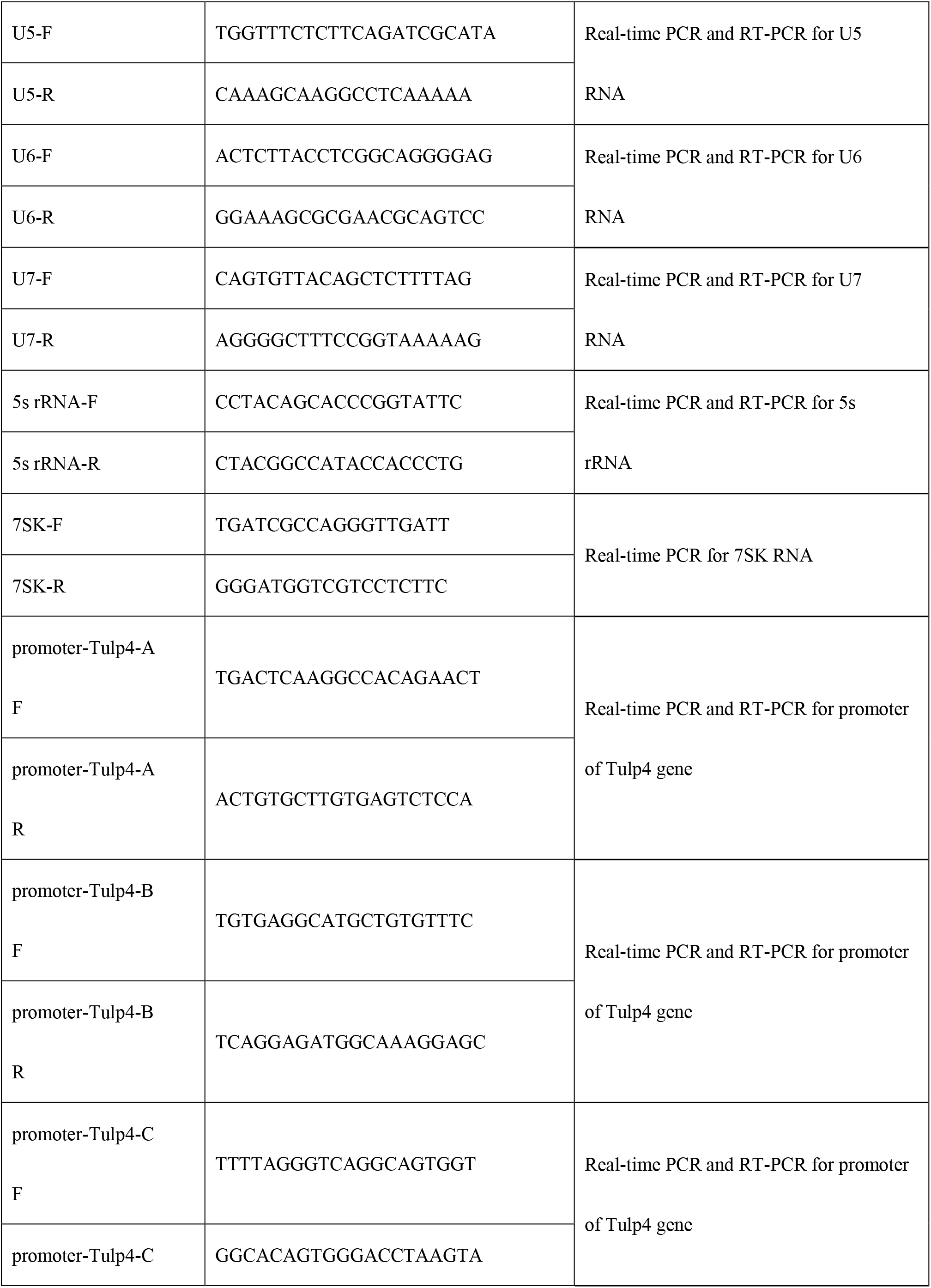
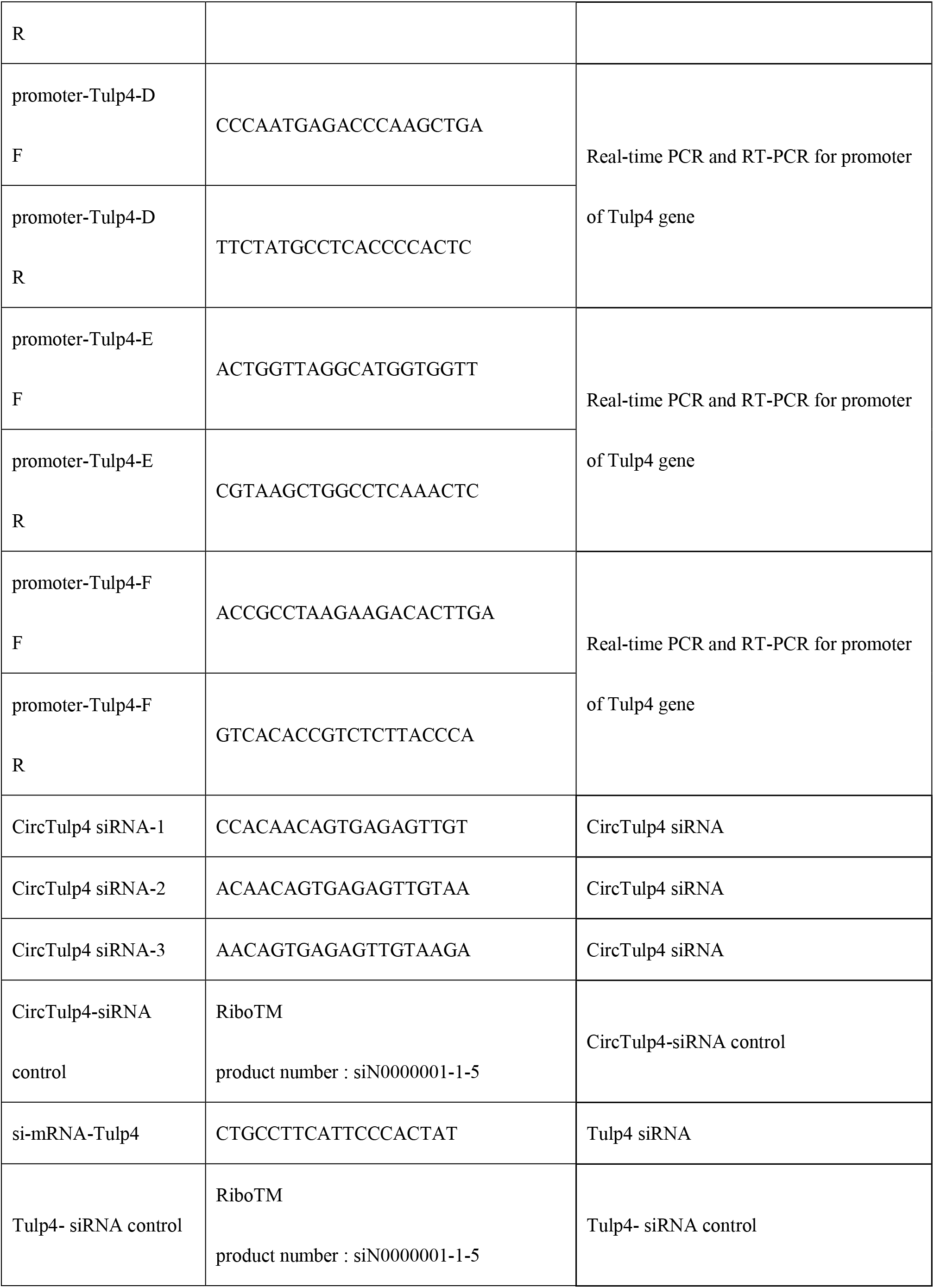
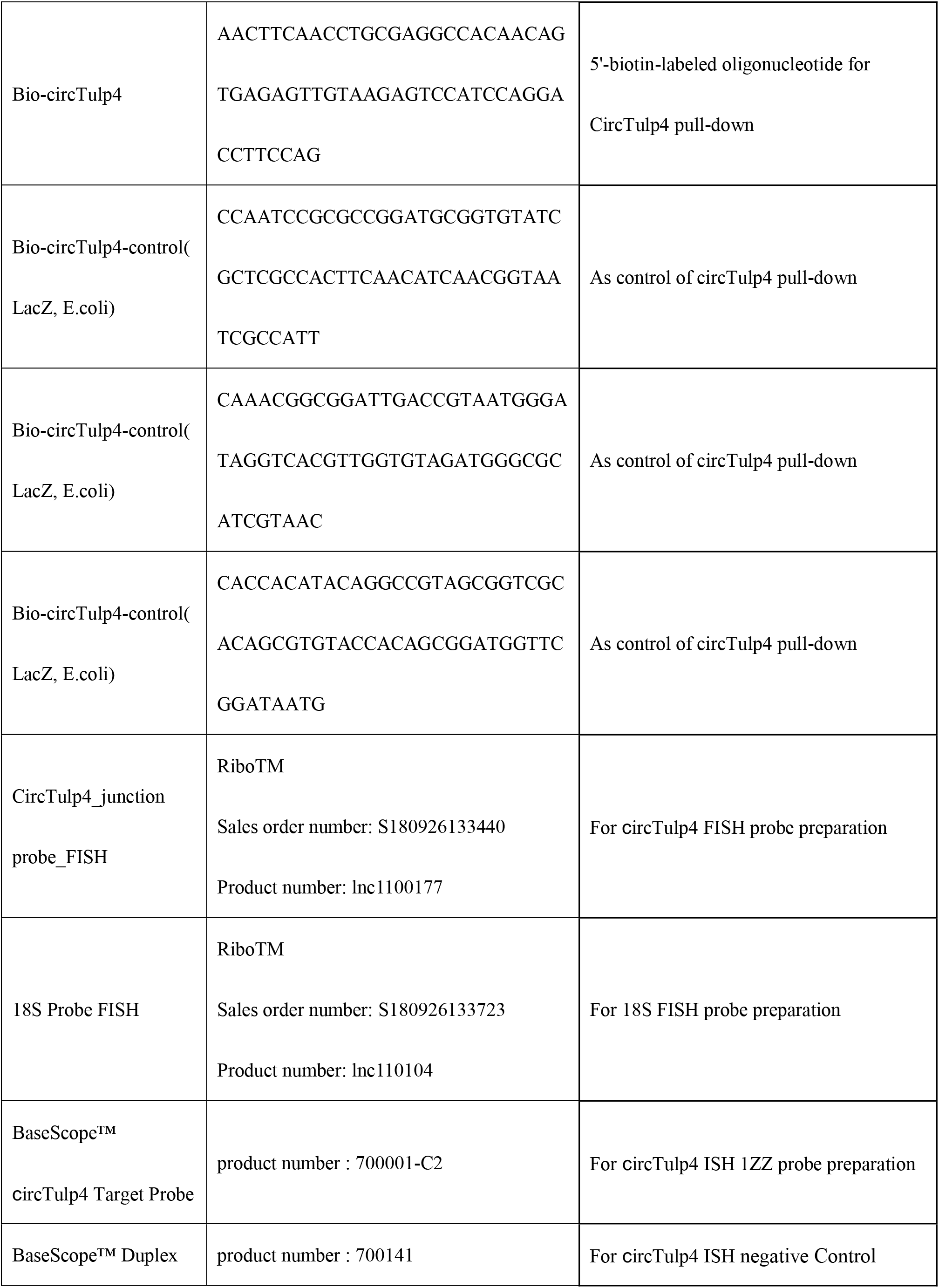
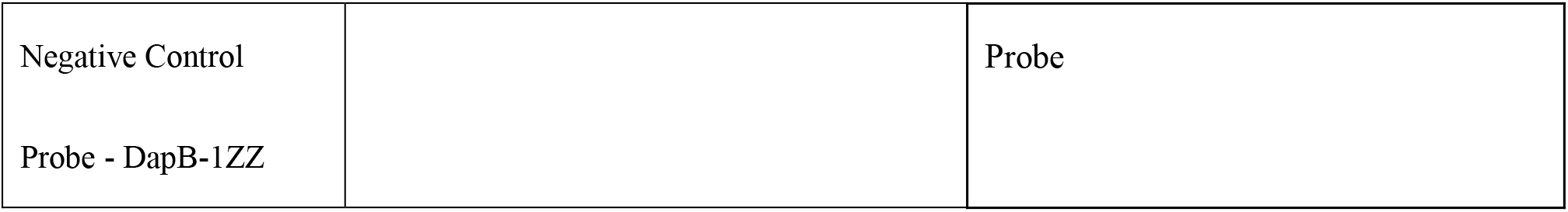
Oligos used in the study.

